# Fbxl10/Kdm2b is required for Kmt2b/Mll2 binding across the genome and regulates H3K4 methylation on bivalent promoters

**DOI:** 10.1101/2025.05.15.654384

**Authors:** Fei Ji, Sharmistha Kundu, Anthony Anselmo, Alexander Morris, Anthony Ducasse, Ayla Ergun, Robert E. Kingston, Ruslan I. Sadreyev

**Affiliations:** Department of Molecular Biology, Massachusetts General Hospital, Boston, MA; Department of Genetics, Harvard Medical School, Boston, MA; Department of Pathology, Massachusetts General Hospital and Harvard Medical School, Boston, MA

**Author notes:** These authors contributed equally to this work (Fei Ji, Sharmistha Kundu, Anthony Anselmo).

## Abstract

The presence of histone modifications associated with both transcriptional repression (H3K27me3) and activation (H3K4me3) on key developmental promoters in embryonic stem cells results from the co-localization of repressive Polycomb group (PcG) and activating Trithorax group (TrxG) protein complexes. Functional interactions between PcG and TrxG on these promoters are not fully understood. Here we focus on the relationships between Fbxl10/Kdm2b, a component of a PcG complex PRC1, and Kmt2b/Mll2, an essential component of TrxG at bivalent promoters. Computational analysis of previously published data revealed genome-wide correlation between chromatin occupancies of these two proteins, suggesting potential crosstalk between Kdm2b and Mll2 at both active and repressed promoters. We tested this hypothesis experimentally and found that loss of Kdm2b resulted in depletion of Mll2 at promoters genome-wide, suggesting that Kdm2b is required for Mll2 occupancy at both bivalent and active promoters. Loss of Kdm2b or the core PRC1 component Ring1b also resulted in the reduction of H3K4me3 specifically at bivalent promoters. These findings provide a direct pathway for cooperation between PcG and TrxG at bivalent promoters, suggesting an unexpected modification to the current model of bivalency. In addition, these findings reveal genome-wide role of Kdm2b independent of the full PRC1 complex.

## INTRODUCTION

Many of the promoters that control key regulatory genes involved in differentiation and early embryonic development are classified as “bivalent” [1]. Bivalent promoters are defined as those marked by two seemingly antagonistic histone modifications, H3K4me3 and H3K27me3. These promoters are controlled in part by specialized protein complexes – the repressive Polycomb group (PcG) and activating Trithorax group (TrxG) complexes. The PcG includes two Polycomb repressive complexes, PRC1 which is responsible for chromatin compaction and histone H2AK119 ubiquitination, and PRC2 which is responsible for H3K27 methylation [2-4]. TrxG is responsible for H3K4 methylation and other aspects of genomic regulation, and associates with active gene promoters and enhancers [5-8]. Aberrations in the activity of these complexes are associated with various human diseases [3,4,6,9,10]. The co-localization of these distinctly functioning complexes at the same promoter results in the co-occurrence of histone modifications associated with both transcriptional repression (H3K27me3) and activation (H3K4me3) [1,6,11,12].

The functional significance of bivalency as a distinct chromatin state has been a topic of active debate. Bivalent genes in embryonic stem cells (ESC) are often key developmental regulators and are transcriptionally inactive. Subsets of bivalent genes are activated during differentiation in a cell lineage-specific manner [1,13,14], and their bivalency is concomitantly resolved [1,15]. In addition to ESCs, smaller numbers of bivalent promoters have been reported in multipotent stem cells [16,17] and differentiated cells [15,17], both in cell culture and in vivo [17-19]. The perturbation of chromatin state by ablation or inactivation of PcG components results in premature activation of many bivalent genes [20-23]. A large number of bivalent genes are also misregulated in cancer [6]. However, the mechanisms of establishing, maintaining, and resolving bivalency, and their role in controlling gene expression are not fully understood.

Subsets of bivalent genes are conserved between human [13,14], mouse [15], and zebrafish [24,25], supporting the functional significance of bivalency in these vertebrates. However, bivalency is not universally conserved through evolution. Co-localization of H3K4me3 and H3K27me3 is largely non-existent in *Drosophila* [26,27] and in late embryos of *Xenopus* [28], although these species still employ both PcG and TrxG proteins to control expression of developmental regulators. Further, since the concept of bivalency originated from ChIP-seq experiments on large populations of cultured cells, it could be attributed to averaging over heterogeneous cells that have either H3K4me3 or H3K27me3 at a given promoter [29]. This argument was additionally supported by high-throughput biochemical evidence against coexistence of the two marks on the same histone molecule in human HeLa cells [30] and by the observed inhibition of H3K27 methylation by PRC2 in the presence of H3K4me3 [31]. More recent direct high-resolution studies, however, suggested that H3K4me3 and H3K27me3 often co-exist at the promoter in the same cell [32], and furthermore, on the same nucleosome, albeit most often on distinct histone molecules [33,34]. As a result, the biological relevance of bivalency is still a subject of active discussion.

To address this functional relevance of co-localization between PcG and TrxG complexes at bivalent promoters, we investigated the relationships between PcG and TrxG proteins at these promoters. First, we performed integrative computational analysis of multiple previously published datasets and surveyed correlations between chromatin occupancies of various PcG and TxG subunits in ESCs, which suggested a potential positive crosstalk between Fbxl10/Kdm2b, a key component of non-canonical PRC1, and Kmt2b/Mll2, a methylase member of TrxG that is essential for H3K4 trimethylation at bivalent promoters [35,36]. Next, to test this surprising computational hypothesis, we experimentally characterized the effects caused by the ablation of Kdm2b and Ring1b, the core component of PRC1, on Mll2 and its enzymatic readout H3K4me3. We show that Kdm2b is required for the maintenance of Mll2. Notably, Mll2 occupancy depends on Kdm2b not only at bivalent promoters, but also at Mll2-occupied active promoters genome-wide, which are depleted of full PRC1 complex. We also show that ablation of Ring1b or Kdm2b results in the reduction of H3K4me3 at bivalent promoters. These results provide an insight into genome-wide role of Kdm2b independent of PRC1 complex and suggest a surprising positive cooperation between supposedly counteracting repressive and active chromatin modifiers at bivalent promoters. This crosstalk supports the notion of bivalency as a distinct functional chromatin state that is maintained, at least in part, through PcG-TrxG cooperation.

## RESULTS

### Promoters with highest PcG occupancy comprise an extreme subset of bivalent promoters

To understand if there is a functional interdependency between TrxG and PcG complexes, we first performed a computational survey of previously published data and analyzed patterns of genome-wide co-localization and quantitative correlation between occupancy levels of different protein subunits and histone modifications. For that, we compiled various publicly available ChIP-seq and RNA-seq datasets from multiple previous studies of TrxG and PcG proteins and their readouts, H3K4me3 and H3K27me3 in mouse ESCs [22,35-41].

In contrast to transcriptionally active promoters, which typically show high levels of TrxG components and H3K4me3 and depletion of most PcG components and H3K27me3, bivalent promoters are characterized by the presence of both H3K4me3 and H3K27me3, as well as PcG and TrxG (Fig. 1A, see also additional examples in S1F Fig). To focus on a high-confidence set of bivalent promoters, we first computationally identified transcription start sites (TSS) with substantial levels of both PcG and TrxG. In order to better understand the patterns of PcG and TrxG occupancy at these bivalent promoters in the whole-genome context, we surveyed quantitative relationships between various PcG and TrxG subunits and histone modifications at all TSS across the genome. This computational survey suggested that bivalent promoters have a unique combination of H3K27me3, H3K4me3, PcG, and TrxG levels, and led to the hypothesis that Kdm2B plays a key role in the interface between PcG and TrxG.

**Figure 1.**
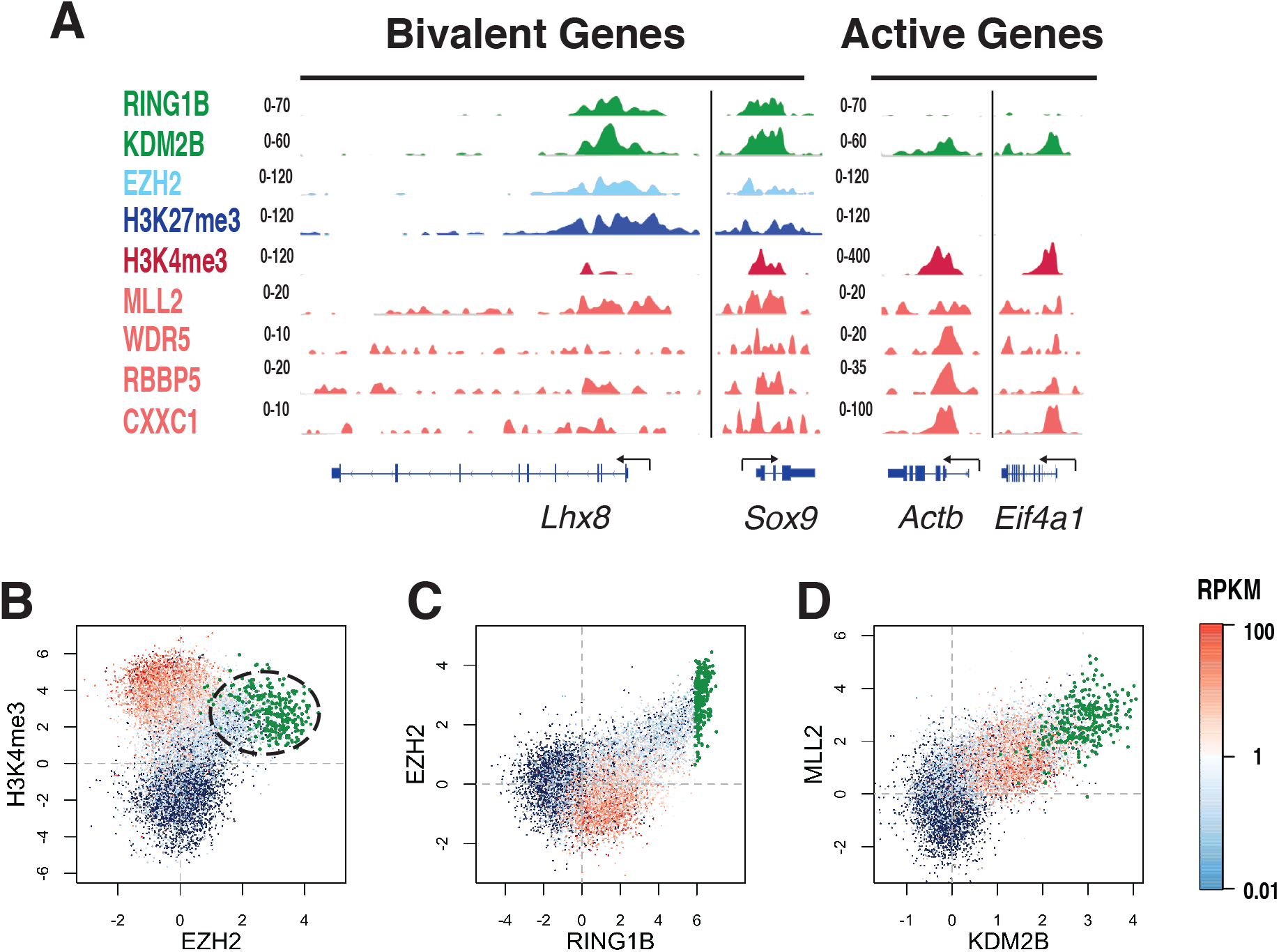
High-PcG promoters have a specific pattern of chromatin marks and chromatin modifiers. A. Examples of ChIP-seq enrichment tracks of H3K4me3, H3K27me3, and members of PcG and TrxG complexes at active and bivalent promoters in wild type ESCs, based on previously published data [22,35-40,71]. B-D: Scatter plots of ChIP-seq tag enrichment in TSS proximal regions (TSS ± 3 Kb) genome-wide. Each point corresponds to an individual TSS, with the level of expression shown by color (from silent as dark blue to highly active as red). TSS with highest Ring1b occupancy are shown as green points. B. Relationship between H3K4me3 and Ezh2 levels on the same TSS involves at least three modes that correspond to three distinct regions in the scatter plot. Bivalent genes occupy a special region of this plot, with intermediate levels of H3K4me3 and highest genome-wide levels of Ezh2. C. At the TSS with a strong enrichment of PRC1 or PRC2, these occupancies are highly correlated. The occupancy of Ezh2, a core enzymatic component of PRC2, is correlated with the occupancy of Ring1b, a core component of PRC1. TSS with high PRC2 occupancy have high PRC1 occupancy, and vice versa. These TSS can be referred to as high-PcG. D. Occupancies of TrxG component MLL2 and PcG component Kdm2b are correlated genome-wide and have high levels at high-PcG promoters.

Our computational analysis confirmed many expected genome-wide correlations between the levels of functionally related components at the same TSS. For example, we observed pronounced quantitative correlations between protein subunits of the same complex (S2, S3E-H, S4B-D Figs) and between complex subunits and the corresponding histone modifications (S1B-D and S2 Figs). Quantitative relationships between TSS-proximal levels of H3K27me3 and H3K4me3, however, have a distinct structure that differs from these simple correlations (S1A Fig). As we described previously [42], all promoters can be roughly divided in three large groups that show different patterns of relationships between H3K4me3 and H3K27me3/PcG (Fig. 1B, S1A Fig). At transcriptionally active promoters (gray and red points), H3K4me3 is high and largely anticorrelated with the levels of H3K27me3, PRC1, and PRC2 (Fig. 1B, S1A,E Fig). At silent promoters (blue points), H3K4me3 is low or absent, and its depletion is correlated with the depletion H3K27me3, PRC1, and PRC2 (Fig. 1B and S1A,E Fig). An important subset of transcriptionally inactive promoters shows enrichment in H3K27me3 and moderate levels of H3K4me3 and is therefore referred to as bivalent [1]. These promoters (marked with a circle in Fig. 1B and S1A Fig) have high occupancies of PRC2 and often PRC1 components (Fig. 1B-D and S1A-E Fig). They occupy the third distinct region of this plot located near the nexus between active and silent branches (Fig. 1B and S1A Fig). This region corresponds to the highest enrichment levels of Ezh2 (> ∼4-fold), Ring1b (> ∼16-fold), and H3K27me3 (> ∼4-fold) combined with a relatively moderate ∼2-10 fold enrichment of H3K4me3 (Fig. 1B and S1A,E Fig).

To study the functional interactions between TrxG and PcG, we further defined a high-confidence subset of most pronounced bivalent promoters. Consistent with previous observations in ESC [11], occupancies of PRC1 and PRC2 are strongly correlated at the promoters where either complex is substantially enriched (Fig. 1C). We therefore refer to these promoters as PcG-occupied. Among PcG-occupied promoters, we further focused on the subset that has the highest enrichment of PRC1 and PRC2 (Fig 1C, green points). We defined this high-PcG subgroup as the top 300 promoters by Ring1b occupancy (Ring1b enrichment >∼60 fold), which represent the most pronounced subset within the bivalent region (Fig. 1B,C and S1A,E Fig) and correspond to the most extreme enrichment of PcG and H3K27me3 combined with moderate levels of H3K4me3. The majority of these promoters control homeobox genes and other early developmental transcription regulators (S1 Table). We used these high-PcG promoter regions as a set of highly confident bivalent promoters and compared the occupancy patterns of various PcG and TrxG components at these promoters to other PcG-occupied promoters and to transcriptionally active promoters with no PcG/H3K27me3 and high TrxG/H3K4me3 densities.

### Genome-wide similarity between chromatin occupancy patterns of Kdm2b and Mll2

Both PcG and TrxG are multisubunit complexes that include various subclasses occurring as distinct subunit combinations. To determine which particular PcG classes or individual subunits co-localize with subunits of TrxG, we computationally analyzed publicly available ChIP-seq data on various PcG and TrxG components in ESCs [22,35-41]. In addition to confirming previously known relationships between these proteins, our analyses revealed unexpected patterns suggestive of previously unknown functional connections (Fig. 1C,D, S2-S4 Figs). Consistent with previous reports in ESCs and other cell types [22,39,40,43-45], we found that in contrast to most PcG components, which mainly occupy bivalent promoters, Kdm2b protein, a member of non-canonical PRC1 [46-48], occupies both bivalent and transcriptionally active promoters (Fig. 1A,D, Fig. 2). Furthermore, we found that TSS-proximal occupancy of Kdm2b is quantitatively correlated with the occupancies of TrxG components Rbbp5, Ash2l, Wdr5, Cxxc1, and Mll2 genome-wide (S2 and S3A-D Figs). Remarkably, these correlations are comparable to the correlations among TrxG components themselves (S2 and S3E-H Figs). This unusual dual association of Kdm2b with TrxG genome-wide (S2 and S3A-D Figs) and with PcG at bivalent genes (S2 and S4A-D Figs) is reminiscent of one specific TrxG component, Mll2 (S2, S3E-H, and S4E-H Figs), which is strongly enriched at active promoters and is also considered the essential H3K4 methylase at bivalent promoters [35,36].

**Figure 2.**
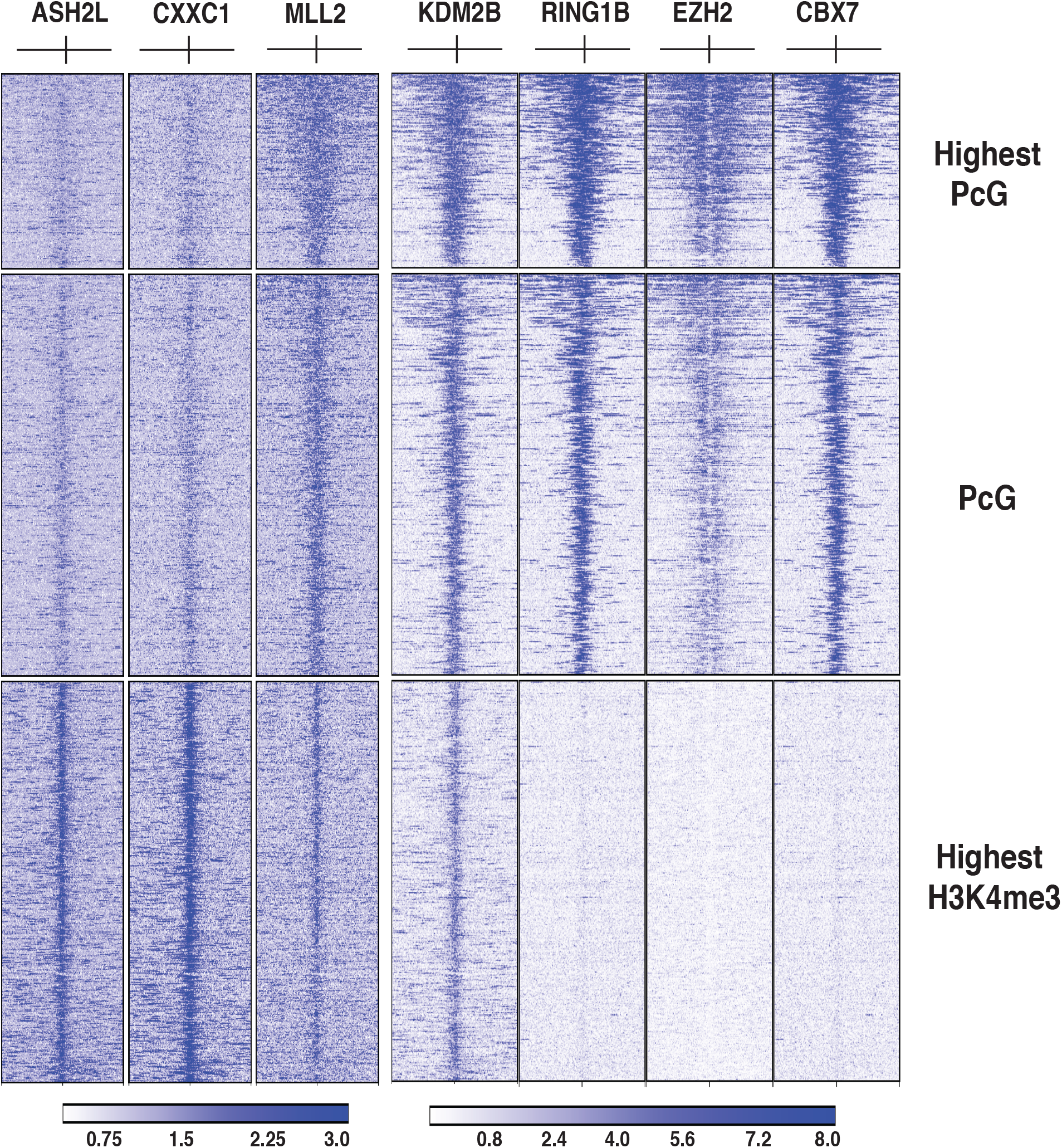
TSS occupancy patterns of MLL2 and KDM2B are similar to each other genome-wide and to PcG and TrxG at high-PcG and active promoters, respectively. ChIP-seq enrichment profiles over individual TSS-proximal regions (TSS ± 10 Kb) for MLL2 [36] and Kdm2b [39] (center) compared to core TrxG components (Ash2l and Cxxc1 [36], left), and PcG proteins (Ring1b [71], Cbx7 [41], and Ezh2 [38], right). The sets of 300 high-PcG TSS, 600 PcG-occupied TSS with lower PcG occupancy, and 600 active TSS with highest H3K4me3 density are shown as separate groups.

This general similarity in the behavior of genomic occupancies between Kdm2b and Mll2 extends to higher quantitative and positional resolution. Since at the level of average enrichment over a TSS-proximal region, the patterns of Kdm2b relationships with TrxG (S3A-D Fig) and PcG proteins (S4A-D Fig) are similar to the patterns for Mll2 (S3E-H and S4E-H Figs), we further compared the detailed occupancy profiles of Kdm2b [39] and Mll2 [36] at individual TSS-proximal regions to the core components of TrxG (Ash2l [36], Cxxc1 [36]) and PcG (Ring1b [39] as a core component of PRC1, Cbx7 [41] as a component of canonical PRC1, and Ezh2 [38] as an enzymatic component of PRC2).

Figure 2 shows these comparisons across three classes of promoters: high-PcG promoters (Fig. 2, top section), other bivalent promoters with more moderate levels of PcG (Fig. 2, middle section), and transcriptionally active promoters with highest levels of H3K4me3 and TrxG combined with depletion of PcG (Fig. 2, bottom section). At high-PcG promoters, all PcG components and Mll2 show a similar pattern of extremely high enrichment across a wide TSS-proximal region spanning many kilobases (Fig. 2). At promoters with more moderate PcG levels, this enrichment is weaker and occurs within narrower regions for both the PcG components and Mll2 (Fig. 2, see also S4 Fig). At active promoters, most PcG proteins, with the notable exception of Kdm2b, are depleted, whereas Mll2 is enriched and localized similar to Kdm2b. Occupancy patterns of other TrxG components Ash2l and Cxxc1 are different: they show a strong enrichment at active promoters and detectable but lower presence at PcG-occupied promoters. Thus, unlike other components, Kdm2b and Mll2 are substantially enriched across all three promoter categories and have similar occupancy patterns (Fig. 2).

Since both Kdm2b and Mll2 bind DNA via their CxxC domain, we addressed the question whether the similarity of their binding patterns is simply due to the presence of CxxC domains and whether other CxxC-containing proteins have similar occupancy patterns. The binding pattern for one of these proteins, the TrxG component Cxxc1 (Cfp1) is shown in Fig. 2 and is different from the patterns of Kdm2b and Mll2. To analyze a more comprehensive set of CxxC-containing proteins, we used publicly available ChIP-seq datasets for Mll4 [49], Kdm2A [39,50], Fbxl19 [51,52], and compared their enrichment patterns to Mll2 [36] and Kdm2b [39] across the same three categories of promoters as shown in Fig. 2. These patterns are shown in S5 Fig. With a possible exception of Fbxl19, neither of these proteins have a strong similarity to Kdm2b and Mll2 in their binding patterns (S5 Fig), suggesting that the similarity between the occupancies of Kdm2b and Mll2 is due to the factors that are more specific than just the presence of a similar DNA binding domain.

In sum, our computational analysis of public datasets suggested that Kdm2b and Mll2 is the only pair of surveyed PcG and TrxG components whose TSS occupancies are quantitatively correlated genome-wide, elevated at high-PcG promoters (Fig. 1D, 2), and show similar patterns of correlation with both PcG and TrxG complexes (Fig. 2, S3 and S4 Figs). These observations suggest a genome-wide association of Kdm2b with a TrxG complex, in particular with Mll2 as a pronounced and universal partner. We hypothesized that this association results from a functional interaction between Kdm2b and Mll2, and this interaction does not require the presence of full PcG complexes, as it is maintained on active promoters in the absence of PcG (Fig. 1A,D, Fig. 2).

### Ablation of Kdm2b results in depletion of Mll2 at all promoters and reduction of H3K4me3 at bivalent promoters

We experimentally tested our hypotheses by analyzing the effect of ablation of PRC1 components on the levels of Mll2 and H3K4me3 at all promoters across the genome, and specifically at high-PcG promoters and active promoters. We compared the effects of (a) ablation of Ring1b, a core component of PRC1 complexes, which would result in the loss of all PRC1 occupancy, and (b) ablation of Kdm2b, which is present only in the subset of non-canonical PRC1 complexes [46-48] but is also present at active promoters (Fig. 1A,D, Fig. 2). We used ChIP-seq to analyze the levels of Mll2 and H3K4me3 at TSS-proximal regions in Kdm2b-null and Ring1b-null ESCs compared with wild-type ESCs. Ring1b [53-56] and Kdm2b deletions [45,57] produce severe developmental defects in mice but do not cause a strong phenotypic effect in cultured ESCs, albeit disrupting their proper differentiation [40].

We were unable to measure Mll2 binding accurately in standard ChIP-seq protocols using commercially available antibodies to mouse Mll2 due to high levels of background noise. To increase signal-to-noise ratio of Mll2 antibodies, we applied whole-genome amplification (WGA) of ChIP material [58-60]. As a more comprehensive approach compared to the microarray-based analyses used in previous studies [60], we subjected the resulting amplified DNA to next-generation sequencing. As a control, applying WGA to low amounts of ChIP material produced with a high-quality H3K4me3 antibody, we observed a good correlation between ChIP-seq signal in WGA-enriched samples and H3K4me3 samples processed with standard ChIP-seq (S6A,B Figs). This correlation largely held across the whole range of GC content, H3K4me3 and PcG density, and promoter activity (S6A,B Figs). When applied to the Mll2 ChIP-seq material, WGA produced the pattern of genome-wide enrichment (Fig. 3A) that was consistent with the known pattern of Mll2 binding. The quantitative Mll2 ChIP-seq enrichment across all promoters showed distinct correlation between our WGA-based experiments and the previously published ChIP-seq data for the GFP-tagged Mll2 in mouse ESCs [36], despite different ChIP approaches and antibodies (S6C Fig). As a result, the sets of Mll2 targets, defined by the promoter regions with enriched ChIP-seq signal, were largely consistent between these two experiments (S6D Fig).

**Figure 3.**
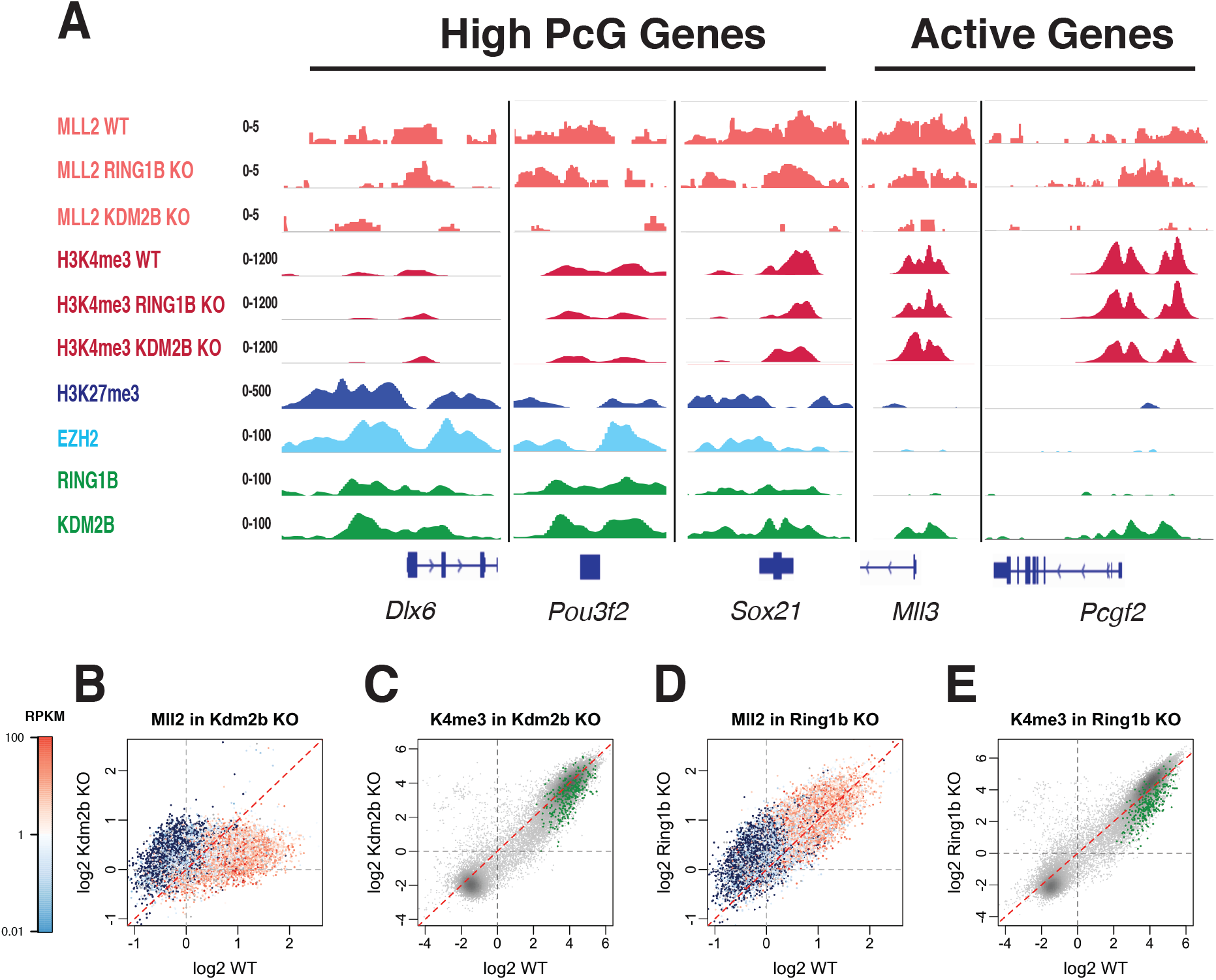
Effects of Kdm2b and Ring1b ablation on the levels of Mll2 and H3K4me3 at individual promoters. A. Examples of ChIP-seq enrichment tracks of MLL2 and H3K4me3 in wild-type, Kdm2b-null, and Ring1b-null ESCs at the TSS of high-PcG and active genes, shown together with the tracks of Ring1b, Kdm2b, and Ezh2 in wild-type ESCs. B-E: Scatter plots of ChIP-seq tag enrichment in all TSS-proximal regions (TSS ± 3 Kb) in mutant vs wild-type ESCs. Each point corresponds to an individual TSS. B. Ablation of Kdm2b results in a strong genome-wide depletion of Mll2. The scatter plot of Mll2 enrichment in Kdm2b-null vs wild type is strongly tilted below diagonal in the area of corresponding to promoters with substantial Mll2 enrichment in wild type. Wild-type gene expression level is shown by the color of the point (from silent as dark blue to highly active as red). C. Ablation of Kdm2b results in a reduction of H3K4me3 at the high-PcG promoters. Scatter plot of H3K4me3 enrichment in Kdm2b-null vs wild type, with the high-PcG promoters marked green. D. Ablation of Ring1b does not have as strong effect on Mll2 occupancy as ablation of Kdm2b. The scatter plot of Mll2 enrichment in Ring1b-null vs wild type is close to diagonal. Wild-type gene expression level is shown by color. E. Ablation of Ring1b results in a reduction of H3K4me3 at the high-PcG promoters. Scatter plot of H3K4me3 enrichment in Ring1b-null vs wild type, with high-PcG promoters marked green.

Compared to wild-type ESCs, Mll2 occupancy was strongly reduced in Kdm2b-null ESCs on TSS across the genome, but not as strongly in Ring1b-null ESCs. At the same time, H3K4me3 enrichment was reduced specifically at PcG-occupied promoters in both mutants, consistent among three biological replicates. Active promoters in wild-type ESCs (e.g. Mll3 and Pcgf2, Fig. 3A) were strongly occupied by TrxG components including Mll2 and were devoid of PcG components, except for Kdm2b, which was substantially enriched at these promoters. PcG-occupied promoters in wild-type ESCs (e.g. Dlx6, Pou3f2, and Sox21, Fig. 3A) were strongly occupied by PRC1 and PRC2 but had a low occupancy of TrxG, except for Mll2 whose substantial enrichment was similar to Kdm2b and Ring1b (Fig. 3A). Ablation of Kdm2b resulted in the reduction of Mll2 occupancy at both active and PcG-occupied promoters (Fig. 3A,B), consistent with the correlation between occupancies of Kdm2b and Mll2 in wild type cells. Ablation of Ring1b, however, did not produce such a dramatic effect on Mll2 occupancy, even at PcG-occupied promoters where Mll2 is co-localized with PRC1. This contrast between the effects of Kdm2b and Ring1b ablation could be observed at both individual promoters (Fig. 3A) and at the genome-wide level of Mll2 occupancies across all promoters (Fig. 3B vs Fig. 3D; see also Fig. 4B vs Fig. 4C).

**Figure 4.**
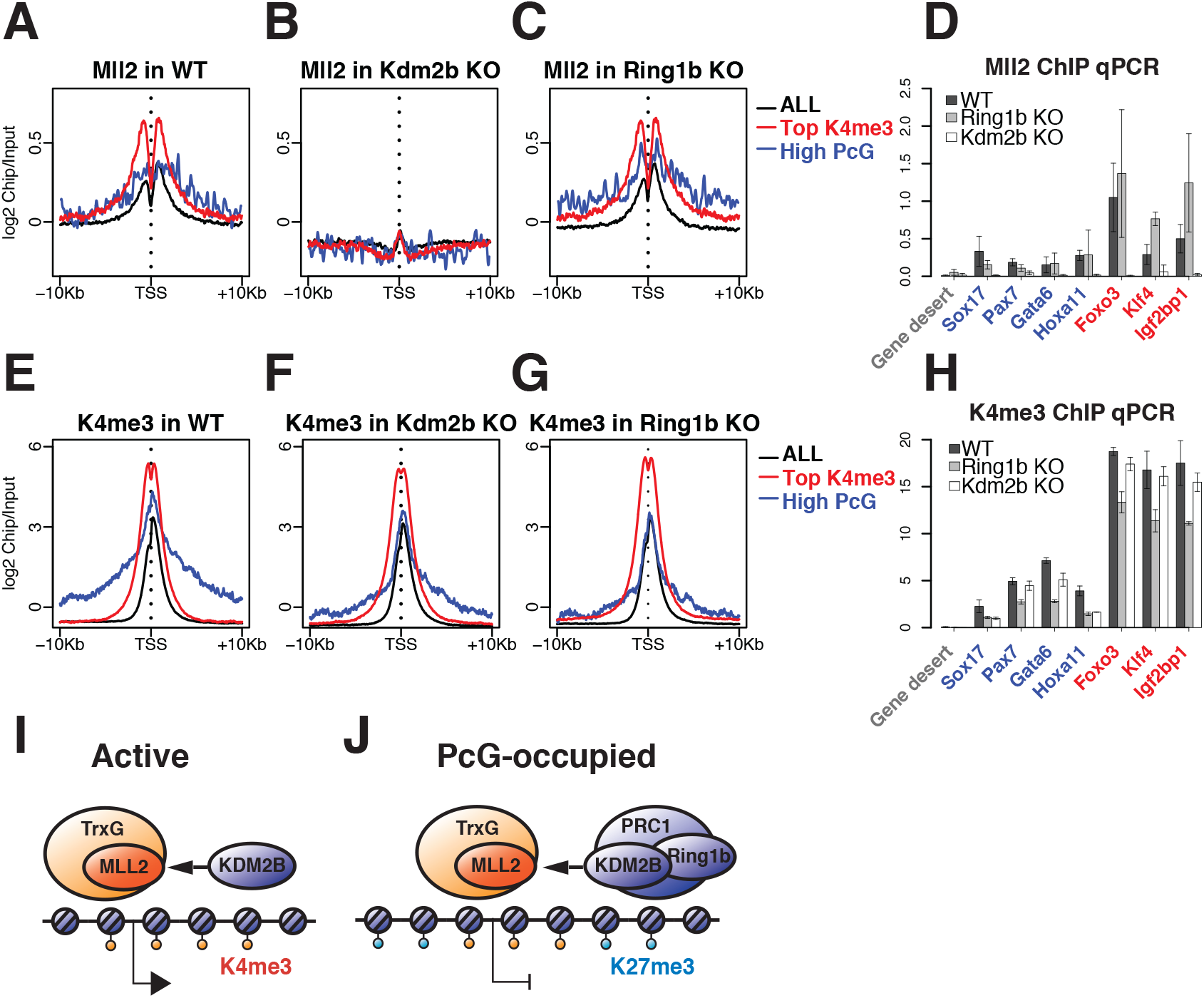
Effects of Kdm2b and Ring1b ablation on the levels of Mll2 and H3K4me3 at different promoter categories. A-C: Average TSS-proximal profiles of MLL2 enrichment for all TSS (black), active TSS with highest H3K4me3 density (red), and high-PcG TSS (blue). A. Wild-type ESCs. B. Kdm2b-null ESCs. C. Ring1b-null ESCs. D. Results of ChIP qPCR analyses of MLL2 enrichment (% input) on TSS of bivalent (Sox17, Pax7, Gata6, Hoxa11) and active genes (Foxo3, Klf4, Igf2bp1) in wild-type, Ring1b-null, and Kdm2b-null ESCs. Gene desert region is shown as a control. Error bars denote SD (n=3). E-G: Average TSS-proximal profiles of H3K4me3 enrichment for all TSS (black), active TSS with highest H3K4me3 density (red), and high-PcG TSS (blue). E. Wild-type ESCs. F. Kdm2b-null ESCs. G. Ring1b-null ESCs. H. Results of ChIP qPCR analyses of H3K4me3 enrichment (% input) on TSS of high-PcG (Sox17, Pax7, Gata6, Hoxa11) and active genes (Foxo3, Klf4, Igf2bp1) in wild-type, Ring1b-null, and Kdm2b-null ESCs. Gene desert region is shown as a control. Error bars denote SD (n=3). I-J: The model of Kdm2b and Mll2 association at active and PcG-occupied promoters. I. At active promoters, Kdm2b binds chromatin in the absence of larger PRC1 and is associated with Mll2 as a part of TrxG. J. At PcG-occupied promoters, Kdm2b association with Mll2 occurs in the presence of PRC1, with Ring1b as its core component. At these promoters, as opposed to active promoters, Mll2 is an essential H3K4me3 methylase whose loss results in the depletion of H3K4me3.

Given the previous reports that the deletion or knockdown of Mll2 reduces the level of H3K4me3 exclusively at bivalent promoters [35,36], we hypothesized that the ablation of Kdm2b and the resulting depletion of Mll2 would similarly reduce H3K4me3 at bivalent promoters. Indeed, Kdm2b-null cells showed reduced enrichment of H3K4me3 at PcG-occupied promoters but not at active promoters (Fig. 3A).

Since the loss of Ring1b did not have such a strong effect on Mll2 occupancy, we expected that ablating Ring1b would impact H3K4me3 less than ablating Kdm2b. Unexpectedly, we found that Ring1b deletion resulted in a similar reduction of H3K4me3 levels specifically at PcG-occupied promoters (Fig. 3A). Loss of either Kdm2b or Ring1b did not result in dramatic changes of high H3K4me3 peaks at active promoters (e.g. Mll3 and Pcgf2 promoters, Fig. 3A), whereas the levels of H3K4me3 at PcG-occupied promoters (e.g. Dlx6, Pou3f2, and Sox21 promoters, Fig. 3A) were reduced in both mutants. As verified by the expression of major ESC marker genes in the RNA-seq experiments and by the alkaline phosphatase staining, neither Kdm2b nor Ring1b ablation caused EScs to enter differentiation, excluding the possibility of cell differentiation being a factor in these effects.

The sharp contrast between the effects of Kdm2b and Ring1b ablation on Mll2 occupancy was apparent when we inspected all individual promoters across the genome. Scatter plot analysis of Mll2 enrichment at individual promoters in mutant vs wild-type ESCs showed that Mll2 levels underwent a strong concerted depletion in Kdm2b-null cells (Wilcoxon *P*-value of 3.5×10^−22^, Fig. 3B). Although the scatter plot in Fig. 3B includes a number of points above diagonal that correspond to a nominal increase of the Mll2 enrichment signal, the vast majority of these points correspond to the sub-threshold values of ChIP to input ratio below 2-fold in both WT or KO that are likely within the level of technical noise. This observed depletion of Mll2 at Mll2 target promoters was largely consistent among multiple biological replicates, as confirmed by pairwise comparisons of four ChIP-seq replicates in wild type cells to three replicates in Kdm2b-null cells (S7 Fig). In wild-type ESCs, Mll2 occupancy generally correlated with gene expression: low or absent Mll2 enrichment largely associated with low or no expression, whereas stronger Mll2 enrichment often associated with higher expression (Fig. 3B, expression levels marked by color). This correlation was eliminated in the Kdm2b-null mutant where Mll2 binding was reduced across the vast majority of its targets regardless of expression (Fig. 3B). This depletion of Mll2 was accompanied by a consistent 2-3-fold reduction of H3K4me3 enrichment at high-PcG promoters (Fig. 3C, green points), which was significantly different from other promoters genome-wide (*P*-value = 1.5×10^−18^, S9A Fig). This 2-3 fold reduction of H3K4me3 levels at bivalent promoters was reminiscent of the previously reported significant but not complete reduction of H3K4me3 at these ESC promoters upon Mll2 ablation [35,36]. The remaining levels of H3K4me3 were likely due to the activity of other TrxG H3K4 methylases (Mll1 and potentially others) that are co-localized with Mll2 at these promoters.

Notably, the ablation of Ring1b did not have such a strong genome-wide effect on Mll2 occupancy. In Ring1b-null cells, Mll2 was maintained closer to wild-type levels across the genome (Fig. 3D). This pattern was largely consistent among pairwise comparisons of four biological ChIP-seq replicates in wild type cells to three replicates in Ring1b-null cells (S8 Fig). However, similar to Kdm2b-null cells, the level of H3K4me3 at high-PcG promoters underwent a consistent reduction (Fig. 3E, green points) that was significantly different from other promoters (*P*-value = 1.9×10^−67^, S9B Fig). As previously shown by Farcas et al [39], Ring1b is not required for Kdm2b recruitment, and ablation of Ring1b in ESCs does not lead to the depletion of Kdm2b at PcG-occupied promoters. Combined with this observation, our results (Fig. 3D) suggest that in Ring1b-null ESCs, where both Kdm2b and Mll2 levels are maintained near wild-type levels, the reduction of H3K4me3 at PcG-occupied promoters (Fig. 3E) may be mediated via an alternative Kdm2b- and Mll2-independent mechanism.

These genome-wide patterns of Mll2 and H3K4me3 enrichment were also observed among promoter categories at higher positional resolution, as shown in Fig. 4A-C,E-G by TSS-proximal positional profiles averaged across all promoters (black lines), the set of 300 high-PcG promoters (blue lines), and the set of 300 active promoters with highest levels of H3K4me3 and no PcG (red lines). Compared to wild-type ESCs (Fig. 4A,E), ablation of Kdm2b resulted in a systematic depletion of Mll2 in every promoter category (Fig. 4B) and a significant 2-3-fold reduction of H3K4me3 specifically at high-PcG promoters (Fig. 4F). Ablation of Ring1b did not have a strong effect on Mll2 occupancy in either promoter category (Fig. 4C) but led to a significant decrease of H3K4me3 enrichment at high-PcG promoters (Fig. 4G). We verified our ChIP-seq analyses by ChIP-qPCR on representative sets of selected high-PcG and active promoters (Fig 4D,H), confirming that Mll2 occupancy was dependent on Kdm2b but did not require Ring1b, and therefore PRC1.

We conclude that the presence of Kdm2b is essential for proper localization and maintenance of Mll2 at both bivalent and active promoters, and that this function of Kdm2b is independent of its role as a member of the PRC1 family of complexes (Fig. 4I,J).

Similar to previous reports [22,61], both Kdm2b-null and Ring1b-null cells showed a consistent reduction of H3K27me3 at high-PcG promoters to lower but still substantial levels (S9C-E Fig, Wilcoxon *P*-values of 2.7×10^−8^ and 9.6×10^−19^, respectively), suggesting a decrease but not complete depletion of PRC2 at these promoters. Reduction of both H3K4me3 and H3K27me3 at a bivalent promoter corresponds to the shift of its chromatin state towards the lower-left corner in the plot of H3K4me3-H3K27me3 relationships (S1A Fig), where promoters are not bound by either PcG or TrxG complexes and are typically silent (S1A Fig). However, in apparent contrast with the observed reduction of H3K4me3 (Fig. 3C,D and 4F,G), our RNA-seq analysis suggested relatively modest but consistent transcriptional upregulation among ∼30% of high-PcG promoters in both Kdm2b-null and Ring1b-null ESCs (S10 Fig). The patterns of this upregulation were similar between two mutants (S10D Fig) and were consistent with previous reports [39,62,63]. In summary, the ablation of either Kdm2b or Ring1b resulted in the reduction of both H3K4me3 and H3K27me3 at high-PcG promoters, accompanied by their partial transcriptional upregulation.

## DISCUSSION

The Integrative computational analysis of public data from multiple previous studies led us to propose and experimentally verify the unexpected hypothesis that a key component of the PcG system interfaces with a key component of the TrxG system to create functional crosstalk. As an enzymatic component of TrxG, Kmt2b/Mll2 is present across the genome at active promoters; it was also shown to be the key enzyme required for H3K4me3 methylation at bivalent promoters [35,36]. Fbxl10/Kdm2b, a key component of non-canonical PRC1 [22,40,46-48] was suggested to occupy and regulate both bivalent and active promoters [22,40,43-45]. Here we demonstrated the unexpected functional association between Kdm2b and Mll2 at promoters across the ESC genome (Fig. 4I,J). Prompted by our initial observation of a correlation between TSS occupancies of these proteins, we found a surprising dependence of Mll2 occupancy on the presence of Kdm2b. The ablation of Kdm2b results in the depletion of Mll2 genome-wide and the reduction of H3K4me3 specifically at bivalent promoters. The ablation of the core PRC1 component Ring1b also leads to the reduction of H3K4me3 at bivalent promoters, but does have a strong effect on Mll2 binding.

These findings support the concept of bivalency as a biologically relevant chromatin state that involves PcG-TrxG cooperation, as opposed to mere coincidence of these complexes at the same promoter. Our results also suggest modifications to the model of bivalency as an intermediate state of equilibrium resulting solely from counteraction between TrxG and PcG complexes. This model would predict an increase of H3K4me3 upon depletion of PcG. The observed reduction of H3K4me3 suggests cooperative interaction of PcG components with TrxG at high-PcG promoters. This cooperation may provide a feedback mechanism for stable maintenance of bivalency and an additional layer of redundant control in preventing spurious activation of key regulator genes. This axis of regulation may be complementary to the recently proposed regulatory mechanism that involves H3K4me2 methylation by another TrxG member, Km2ta/Mll1 as a key factor in recruitment and maintenance of PRC2 [64].

Our results also suggest that Kdm2b, a component of non-canonical PRC1, has a wider function outside of PRC1-occupied promoters. The genome-wide role of Kdm2b in the maintenance of Mll2 at both PcG and non-PcG promoters (Fig. 4I,J) raises the possibility that Kdm2b may be a part of a pioneering complex that primes CpG islands at TSSs at early developmental stages [36]. In this scenario, the role of Kdm2b as a component of non-canonical PRC1 may be a secondary function performed in the context of PcG-occupied promoters (Fig. 4J). Kdm2b binds unmethylated CpG islands through its CXXC domain, which was proposed to be essential for the recruitment of other PRC1 components including Ring1b to CpG rich promoters [22,40]. Whether this CpG binding or H3K36 demethylase activity of Kdm2b are required for its role in the maintenance of Mll2 levels is an interesting topic for further investigation. However, given the strong presence of Kdm2b at both bivalent and non-bivalent loci, the specificity of recruitment and maintenance of PcG at bivalent promoters is likely to be controlled by factors other than Kdm2b occupancy, e.g. by Mll1-mediated H3K4me2 methylation as a recently proposed candidate [64].

The ablation of Ring1b, a core PRC1 component, results in the loss of PRC1 from its chromatin targets. Ring1b ablation, however, did not have as strong effect on Mll2 occupancy as the ablation of Kdm2b, which strongly depleted Mll2 at both bivalent and active promoters. Thus, although the depletion of Kdm2b has been shown to partially reduce Ring1b occupancy at PcG-occupied promoters [39,40,63], the observed functional association between Kdm2b and Mll2 seems to be independent of Ring1b and PRC1. This independence is consistent with the previously reported independence of Kdm2b occupancy from the presence of Ring1b at PcG target promoters [39]. It is also reminiscent of the recently reported difference between the roles of Ring1b and Kdm2b in protection against ectopic DNA methylation: ablation of Kdm2b leads to de novo DNA methylation at bivalent promoters, whereas ablation of Ring1b does not [45]. Whether this role of Kdm2b in protecting PcG sites against ectopic DNA methylation is connected to Mll2 maintenance can be further addressed by the analysis of DNA methylation at PcG promoters upon the depletion of Mll2.

In Ring1b-null ESCs, the reduction of H3K4me3 at high-PcG promoters (Fig. 3A,E, 4G,H) in the continued presence of Mll2 (Fig. 3A,D, 4C,D) may suggest the existence of additional mechanisms of PRC1-TrxG interaction that are not mediated through Mll2 occupancy. Since the ablation of Ring1b does not have a large effect on the levels of Kdm2b at PcG-occupied promoters [39], the involvement of Kdm2b occupancy in the reduction of H3K4me3 in Ring1b-null ESCs is also unlikely. Instead, PRC1 could affect H3K4me3 levels either by affecting the methylase activity of Mll2 without affecting its binding, or through other enzymatic components of TrxG complex (Mll1,3,4, Set1 proteins). The presence and activity of these non-Mll2 components at bivalent TSS are consistent with the reduced but detectable presence of H3K4me3 observed in ESCs with MLL2 knockout [36] and knockdown [35]. The activity of one of these non-Mll2 components, Mll1, albeit as a H3K4me2 methylase, can be essential for maintaining promoter bivalency [64].

Paradoxically, the ablation of PcG components and the resulting transcriptional upregulation of many bivalent genes coincided with the reduction of active histone modification H3K4me3 at these genes. This may suggest that the function of Mll2 and H3K4me3 at bivalent promoters is not necessary for transcriptional activation, and that the observed upregulation may be driven by other factors and chromatin modifications. The depletion of Mll2 and H3K4me3 upon the removal of PcG components also supports the hypothesis of a specialized mechanism for the targeted activation of bivalent promoters during normal cell differentiation, similar to the model proposed by Denissov et al [36]. In addition to the release of PcG-mediated repression, this mechanism requires active recruitment of other H3K4 methylases, with Set1 as a likely candidate, which would increase the level of H3K4me3 and contribute to the transition of the bivalent promoter to a fully active chromatin state. Finally, the observed reduction of H3K4me3 at upregulated promoters in ESCs raises the possibility that in human diseases associated with the loss of PRC1 function, epigenetic states of upregulated developmental genes may be different from the fully resolved active state in differentiated cells, which warrants a more detailed investigation of these states and approaches to their correction.

## MATERIALS AND METHODS

### Cell culture

Wild type CJ7 and mutant ESC lines were maintained on mitomycin-C inactivated MEF feeder layers (EMD Millipore). Mutant ESC lines used in this study: *Ring1b*^-/-^ (Eskeland et al., 2010), and *Kdm2b*^*T/T*^ (Boulard et al., 2015). These were generous gifts from Wendy Bickmore, and Timothy Bestor labs. All ESC were cultured in DMEM supplemented with 15% fetal bovine serum (Hyclone), Penicillin/Streptomycin, Glutamax, non-essential amino acids, and 1000U/ml of leukemia inhibitory factor (EMD Millipore). All ESC were depleted of feeder cells prior to use in experiments.

### Chromatin immunoprecipitation (ChIP)

Cells were crosslinked for 10 minutes in 1% formaldehyde at room temperature and quenched with 125mM Glycine. ChIP was performed as previously described (Carey et al., 2009), with minor modifications. Antibodies used: H3K4me3 (Millipore, 04-745), H3K27me3 (Cell Signaling Technology, C36B11), MLL2 (Santa Cruz, sc-68678). For MLL2 ChIP, isolated ChIP DNA was processed using a WGA2 kit (Sigma) according to manufacturer’s specifications. All ChIP-seq experiments were performed in 3 or more biological replicates. Some of these samples were further processed for high throughput sequencing.

### ChIP-sequencing and analysis

Sequencing was performed on Illumina HiSeq 2500 instrument. H3K4me3 and H3K27me3 samples were sequenced to approximately 30 million of paired-end 50 bp reads per sample. WGA-enriched MLL2 samples were sequenced to approximately 30 million of paired-end 100 bp reads per sample, followed by computational trimming of primer sequences. Reads were aligned against the mm9 reference genome using BWA [65] with mem option. Alignments were filtered for uniquely mapped reads and alignment duplicates were removed. Enrichment over TSS-proximal regions (TSS ± 3 Kb) was calculated as the ratio of ChIP and input read counts followed by mean shifting. TSS-proximal positional profiles were calculated using standalone version of deepTools [66]. Input-normalized coverage tracks were generated using SPP [67].The same workflow was used for the analysis of publicly available mouse ESC ChIP-seq datasets for H3K4me3, H3K27me3, Kdm2b, Ring1b, Cbx7, Ezh2, Suz12, Rbbp5, Wdr5, Ash2l, Cxxc1, and Mll2 (GEO accession numbers GSE36905, GSE40860, GSE37930, GSE41267, GSE42466, GSE55698, GSE48172, GSE52071, GSE22934).

### RNA Analysis

Total RNA was isolated from two biological replicates of each cell type using TRIzol (Invitrogen), and DNase treated. Sequencing libraries were prepared using polyA-selection followed by NEBNext Ultra Directional RNA library preparation protocol (New England Biolabs). Sequencing was performed on Illumina HiSeq 2500 instrument, resulting in approximately 30 million of 50 bp reads per sample. Sequencing reads were mapped to the mouse transcriptome (mm9 RefSeq annotation) using STAR [68]. Gene expression counts were calculated using HTSeq v.0.6.0 [69]. Calculation of expression values and differential expression analysis were performed using EgdeR package [70].

## ACKNOWLEDGEMENTS

We thank members of the Sadreyev lab for many valuable discussions. This work was supported by the National Institutes of Health (P30 DK040561, R.I.S.).

## AUTHOR CONTRIBUTIONS

RIS, SK, and FJ conceived of the computational analyses and experiments. Computational analyses: FJ, AA, AE. Experiments: SK. Writing – original draft: FJ, SK, AA, and RIS. Writing – review and editing: SK, REK, FJ, and RIS. Funding acquisition: REK and RIS.

## DATA AVAILABILITY

We used publicly available GEO datasets GSE36905, GSE40860, GSE37930, GSE41267, GSE42466, GSE55698, GSE48172, GSE52071, GSE22934. ChIP-seq and RNA-seq datasets produced in this work will be deposited in GEO.

