## Supplemental Figures for "Fbxl10/Kdm2b is required for Kmt2b/Mll2 binding across the genome and regulates H3K4 methylation on bivalent promoters"

#### S1 TABLE (Excel file Table_S1.xlsx)

**Genes with top TSS-proximal occupancy of Ring1b in ESC.** Gene name, mm9 genomic coordinates, and input-normalized ChIP-seq enrichment of Ring1b, Ezh2, Kdm2b, and Mll2 are indicated.

#### SUPPLEMENTARY FIGURE LEGENDS

**S1 Fig. Relationships between TSS-proximal levels of K27me3, Ezh2, and K4me3 genome-wide.**

Scatter plots of ChIP-seq tag enrichment in all TSS-proximal regions (TSS ± 3 Kbp), based on previously published datasets for wild-type ESCs [1-3]. Each point corresponds to an individual TSS, with the wild-type expression level shown by color (from silent as dark blue to highly active as red). High-PcG genes are marked green.

A. H3K27me3 and H3K4me3 have different relationships on active (high H3K4me3, upper area of the plot) and silent genes (depleted H3K4me3, lower area of the plot). Bivalent genes largely occupy a separate area of moderate H3K4me3 and highest H3K27me3 densities. This area corresponds to a high PcG occupancy (see main Figures 1B,C).

B. H3K27me3 density correlates with Ezh2 occupancy.

C. H3K4me3 density largely correlates with Cxxc1 occupancy.

D. H3K4me3 density largely correlates with Mll2 occupancy.

E. Relationship between H3K4me3 and Ring1b levels: bivalent genes occupy a special region of the plot, with high-PcG genes as an extreme subset.

F. Additional examples of ChIP-seq enrichment tracks of H3K4me3, H3K27me3, and members of PcG and TrxG complexes at active and bivalent promoters in wild-type ESCs.

**S2 Fig. Quantitative correlations of Mll2 and Kdm2b occupancies with other TrxG and PcG**

**components on the same TSS, based on previously published data** [1-9].
Heatmap of pairwise Pearson correlation coefficients between TSS-proximal (TSS ± 3 Kbp) levels of ChIP-seq enrichment of K4me3, K27me3, and components of TrxG and PcG complexes on all promoters compared to the same correlations among the subset of high-PcG promoters. A. All promoters. B. High-PcG promoters.

**S3 Fig. Mll2 and Kdm2b occupancies have similar quantitative correlations with TrxG components on TSSs across the genome.** Scatter plots of ChIP-seq tag enrichment (log2 scale) at all TSS-proximal regions (TSS ± 3 Kbp), based on previously published datasets for wild-type ESCs [3-5,7-9]. Each point corresponds to an individual TSS, with the wild-type level of gene expression shown by color (dark blue to red). High-PcG promoters are marked green. A. Kdm2b vs Ash2l. B. Mll2 vs Ash2l. C. Kdm2b vs Rbbp5. D. Mll2 vs Rbbp5. E. Kdm2b vs Wdr5. F. Mll2 vs Wdr5. G. Kdm2b vs Cxxc1. H. Mll2 vs Cxxc1.

**S4 Fig. Mll2 and Kdm2b occupancies have similar quantitative correlations with PcG components**[1-6]**.** Each point corresponds to an individual TSS, with the wild-type expression level shown by color. High-PcG promoters are marked green. A. Kdm2b vs Ring1b. B. Mll2 vs Ring1b. C. Kdm2b vs Cbx7. D. Mll2 vs Cbx7. E. Kdm2b vs Ezh2. F. Mll2 vs Ezh2. G. Kdm2b vs Suz12. H. Mll2 vs Suz12.

**S5 Fig. TSS occupancy patterns of Mll2 and Kdm2b, both of which contain a CXXC DNA-binding domain, are not universally similar to the patterns of other CXXC-containing proteins.**
Enrichment profiles over individual TSS-proximal regions (TSS ± 10 Kbp) were calculated from the published ChIP-seq datasets for Mll2 [3] and Kdm2b [4], and compared to the profiles of other CXXC-containing proteins: Mll4 [10], Kdm2a [4,11] (two independent datasets), Fbxl19 [12], and Tet1 [13,14] (two independent datasets). The sets of 300 high-PcG TSS, 600 PcG-occupied TSS with lower PcG occupancy, and 600 active TSS with highest H3K4me3 density are shown as separate groups.

**S6 Fig. Results of whole-genome amplification (WGA) of H3K4me3 and Mll2 ChIP material were consistent with standard ChIP-seq protocols.**

A. Results of ChIP qPCR analyses of H3K4me3 enrichment (% input) in wild-type ESCs at high-PCG and active promoters, selected promoters with high AT sequence content, and negative control regions with depleted H3K4me3. GD, intergenic region in a gene desert. For each region, three enrichment values are shown for WGA with increasing starting amounts of purified ChIP DNA, starting with low quantities.

B. WGA did not introduce strong systematic biases in H3K4me3 ChIP-seq enrichment values. Scatter plot of TSS-proximal (TSS ± 3 Kbp) ChIP-seq enrichment values for H3K4me3, produced by ChIP followed by WGA vs a standard ChIP protocol. Each point corresponds to an individual TSS, with the level of wild-type expression shown by color.

C. Scatter plot of correlation between the ChIP to input enrichment of Mll2 ChIP-seq signal at TSS-proximal regions across the genome in our ChIP-seq experiments using an Mll2 antibody followed by WGA (y axis) vs previously published ChIP-seq experiment using a GFP antibody in ESCs with GFP-tagged Mll2 [3] (x axis).

D. Venn diagram showing a strong consistency between the sets of target promoters with Mll2 enrichment defined in our ChIP-seq experiments followed by WGA and in the previously published ChIP-seq experiment using a GFP antibody in mouse ESCs with GFP-tagged Mll2 [3].

**S7 Fig. Ablation of Kdm2b results in a strong genome-wide depletion of Mll2 on TSS: consistency between biological replicates of ChIP-seq experiments.** TSS-proximal enrichment of MLL2 is compared for three biological replicates in Kdm2b-null ESCs vs four biological replicates in wild-type ESCs.

**S8 Fig. Ablation of Ring1b does not result in a strong genome-wide depletion of Mll2 on TSS: consistency between biological replicates of ChIP-seq experiments.** TSS-proximal enrichment of Mll2 is compared for three biological replicates in Ring1b-null ESCs vs four biological replicates in wild-type ESCs.

**S9 Fig. A-B: Both Kdm2b and Ring1b ablation resulted in a reduction of H3K4me3 at high-PcG promoters.** Boxplots of the change of H3K4me3 ChIP-seq enrichment (log2 scale) in Kdm2b KO (A) and Ring1b KO (B) compared to wild type among all promoters (gray) and high-PcG pomoters (green). Statistical significance of the difference is indicated by P-value.

**C-E: Kdm2b-null and Ring1b-null ESCs have reduced but still the highest genome-wide levels of H3K27me3 at high-PcG promoters.**

C-D: Scatter plots of H3K27me3 ChIP-seq tag enrichment (log2 scale) at all TSS-proximal regions (TSS ± 3 Kb) across the genome in mutant vs wild-type ESCs. High-PcG promoters are marked green. A. Ring1b-null vs wild type. B. Kdm2b-null vs wild type.

E. Results of ChIP qPCR analyses of K27me3 enrichment (% input) on TSS of bivalent genes (Sox17, Pax7, Gata6, Hoxa11, Hoxa2, Ngn1) in wild-type, Ring1b-null, and Kdm2b-null ESCs. A gene desert region is shown as a control.

**S10 Fig. Ablation of Kdm2b or Ring1b results in the upregulation of expression from many PcG-occupied promoters.**

A. Heatmap of expression values in differentially expressed genes in Ring1b-null and Kdm2b-null mutants vs wild-type ESCs, two replicates.

B. Scatter plot of expression values for all genes in Ring1b-null vs wild type ESCs. Genes with high-PcG promoters are marked green.

C. Scatter plot of expression values of all genes in Kdm2b-null vs wild type ESCs. Genes with high-PcG promoters are marked green.

D. Individual gene expression changes in Ring1b-null and Kdm2b-null cells are quantitatively similar. Scatter plot of expression changes in Ring1b-null compared to wild type (x-axis) vs changes in Kdm2b-null compared to wild type (y-axis).

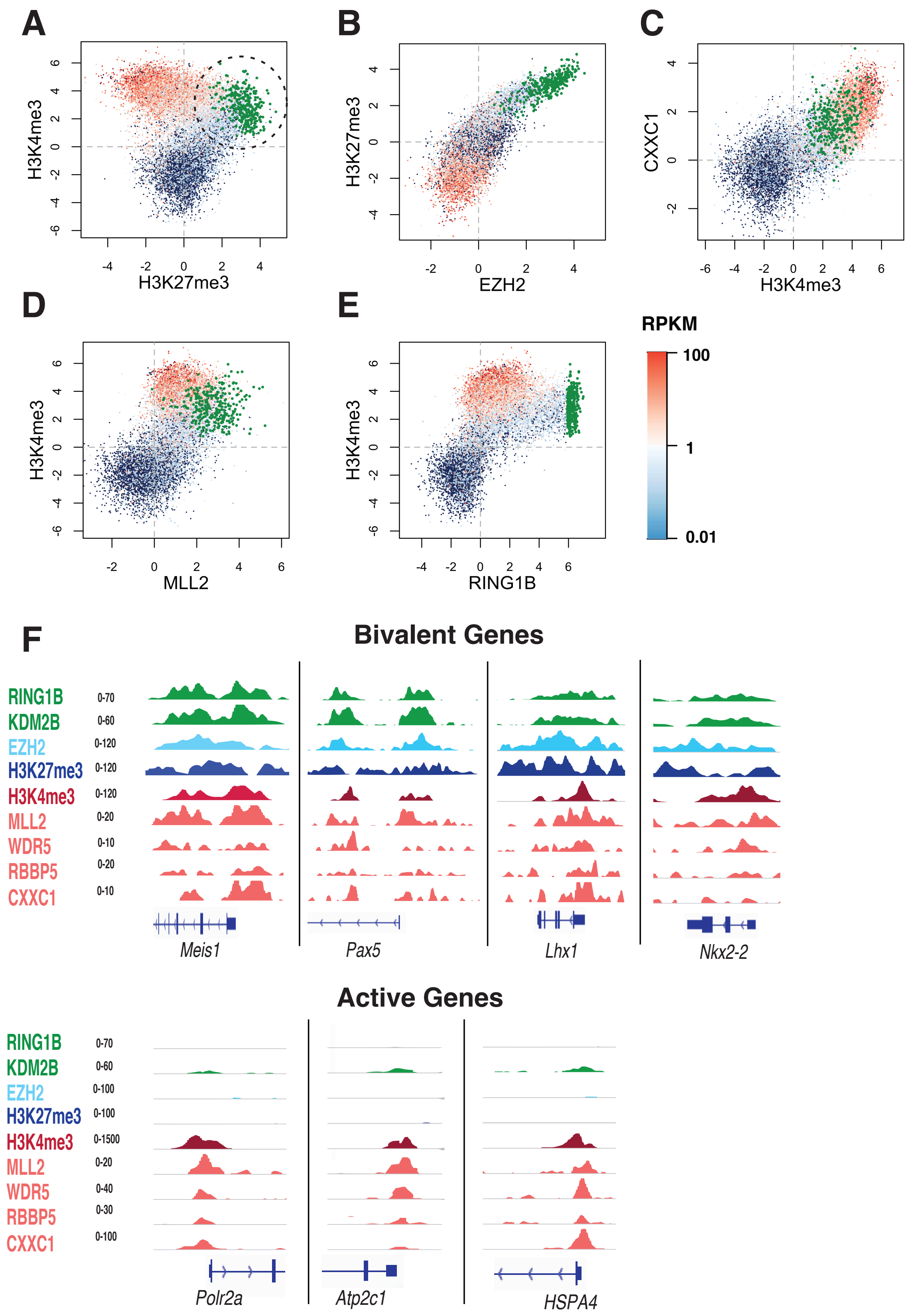
A
B
C
H3K4me3
H3K27me3
H3K27me3
EZH2
CXXC1
H3K4me3
D
E
H3K4me3
MLL2
H3K4me3
RING1B
RPKM
100
1
0.01
F
Bivalent Genes
RING1B
KDM2B
EZH2
H3K27me3
H3K4me3
MLL2
WDR5
RBBP5
CXXC1
0-70
0-60
0-120
0-120
0-120
0-20
0-10
0-20
0-10
Meis1
Pax5
Lhx1
Nkx2-2
Active Genes
RING1B
KDM2B
EZH2
H3K27me3
H3K4me3
MLL2
WDR5
RBBP5
CXXC1
0-70
0-60
0-100
0-100
0-1500
0-20
0-40
0-30
0-100
Polr2a
Atp2c1
HSPA4

#### Ji_FigS2

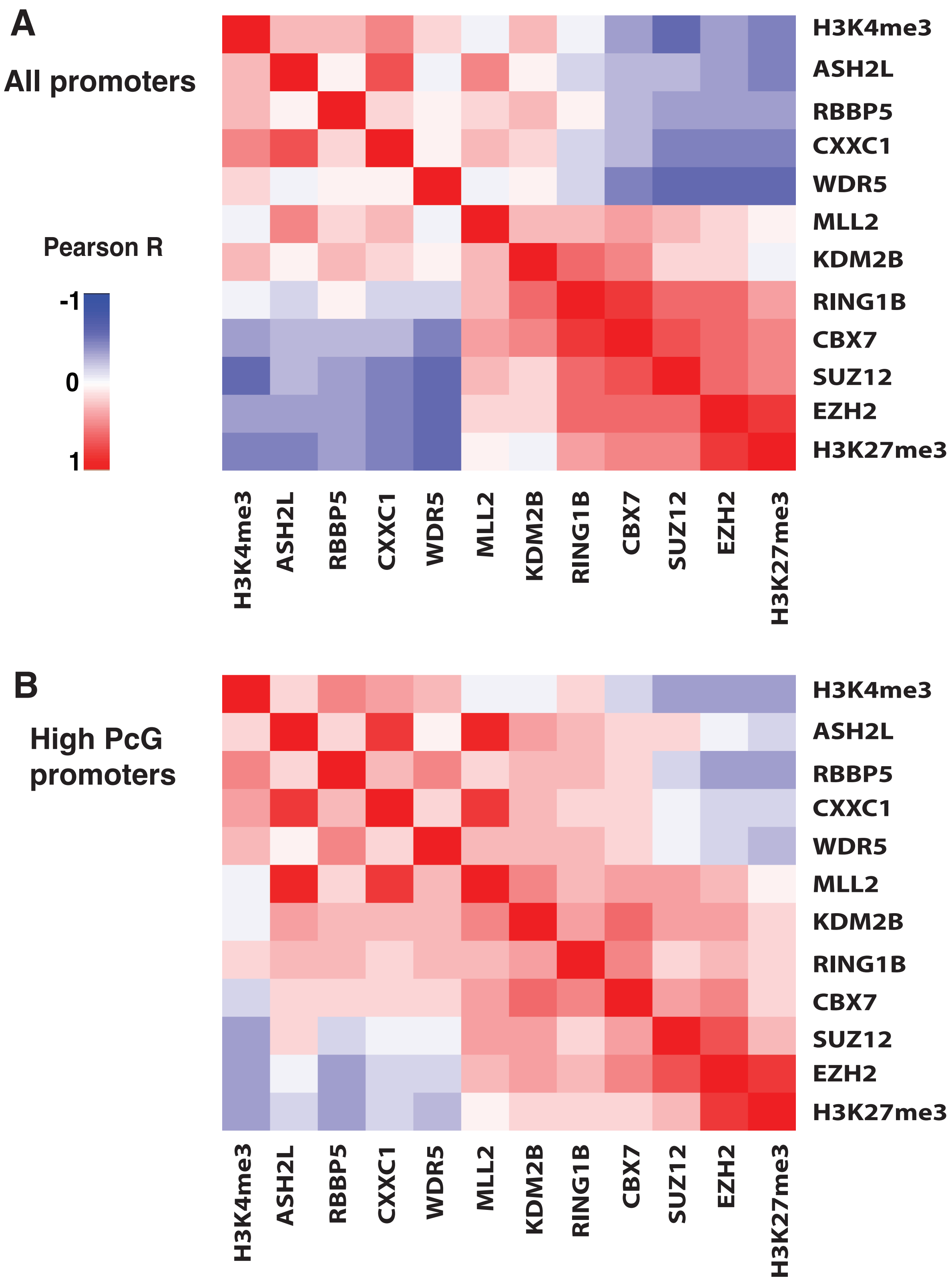

A
All promoters
Pearson R
-1
0
1
H3K4me3
ASH2L
RBBP5
CXXC1
WDR5
MLL2
KDM2B
RING1B
CBX7
SUZ12
EZH2
H3K27me3
B
High PcG promoters

**Ji_FigS3**

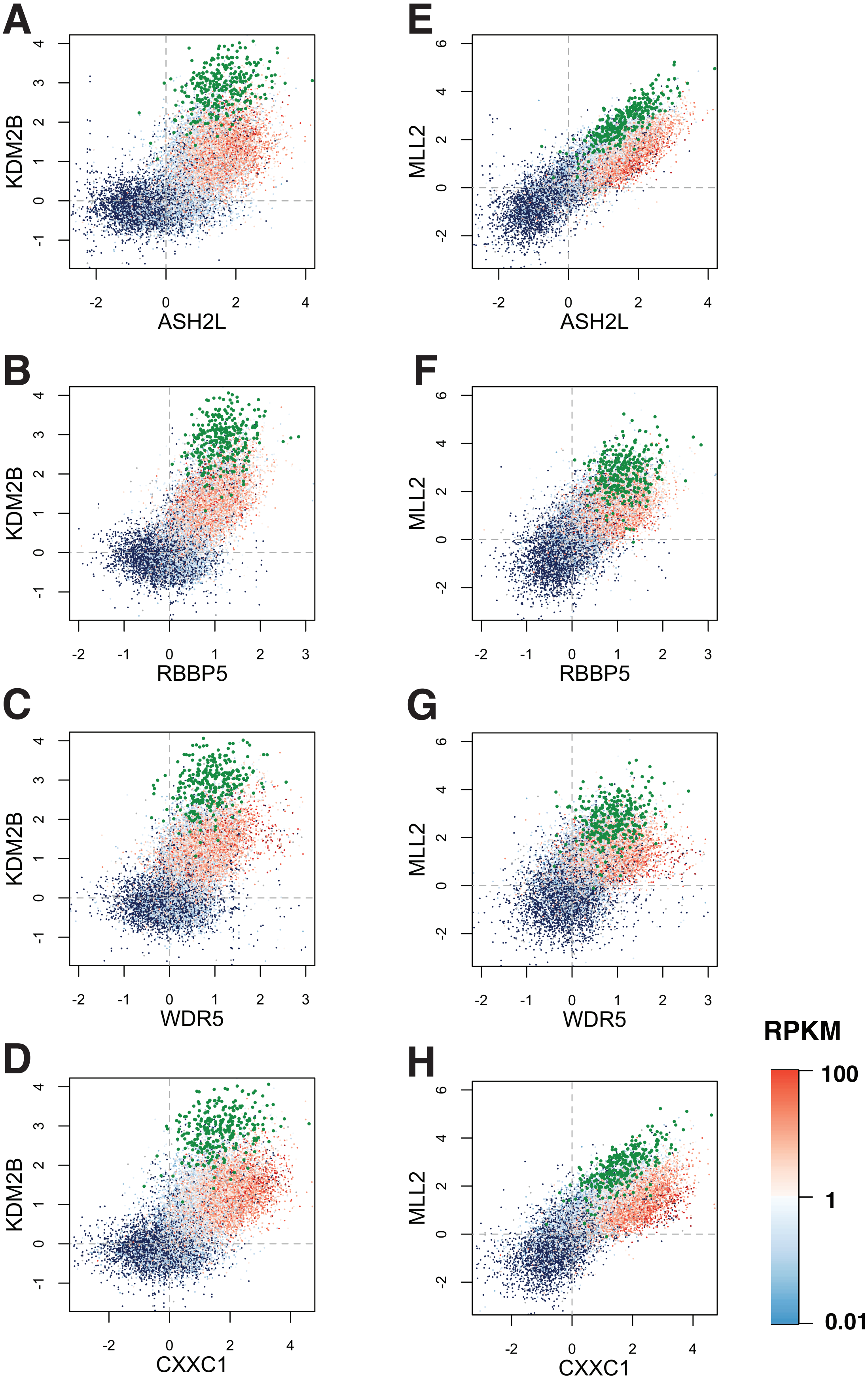

**Ji_FigS4**

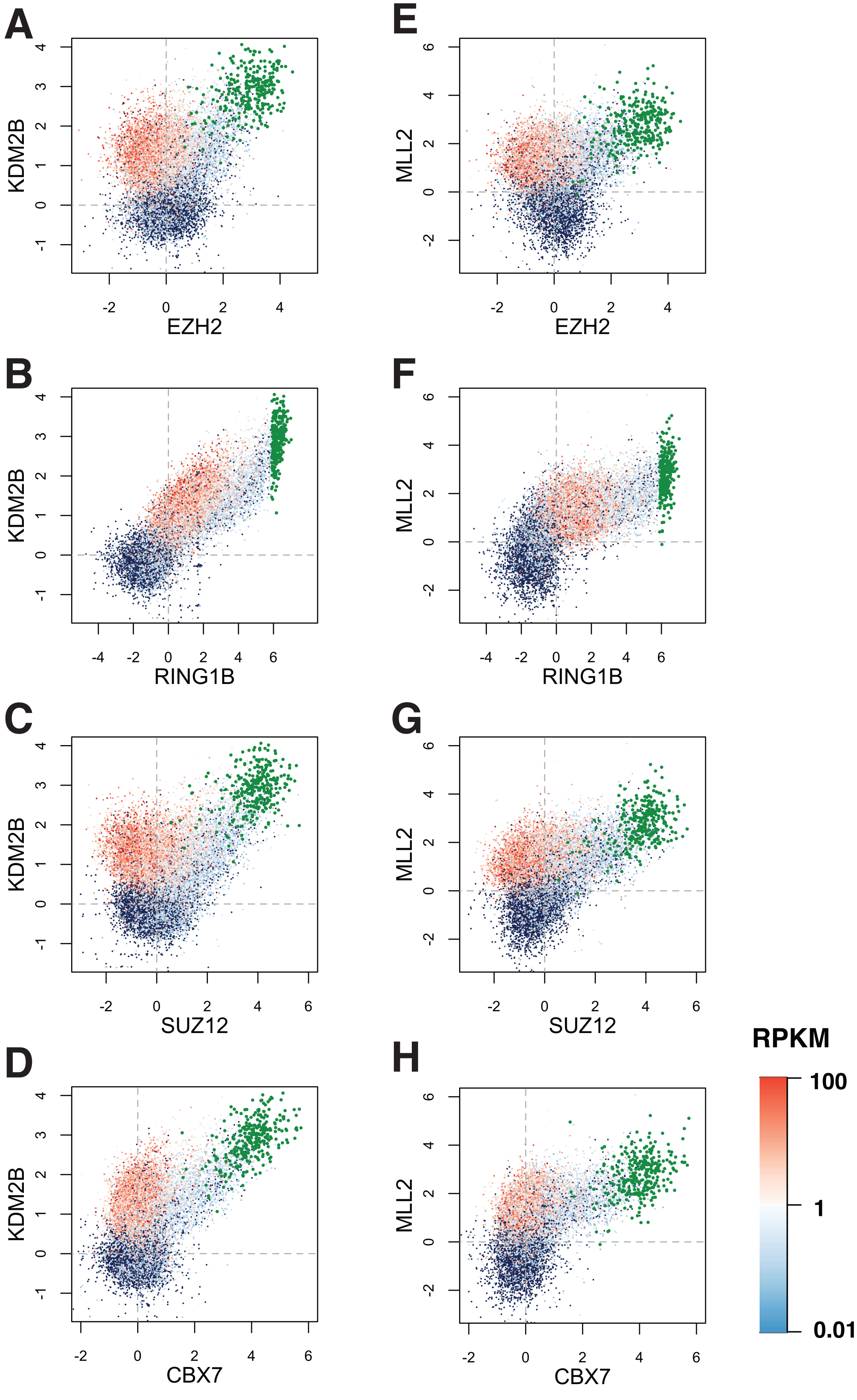

#### Ji_FigS5

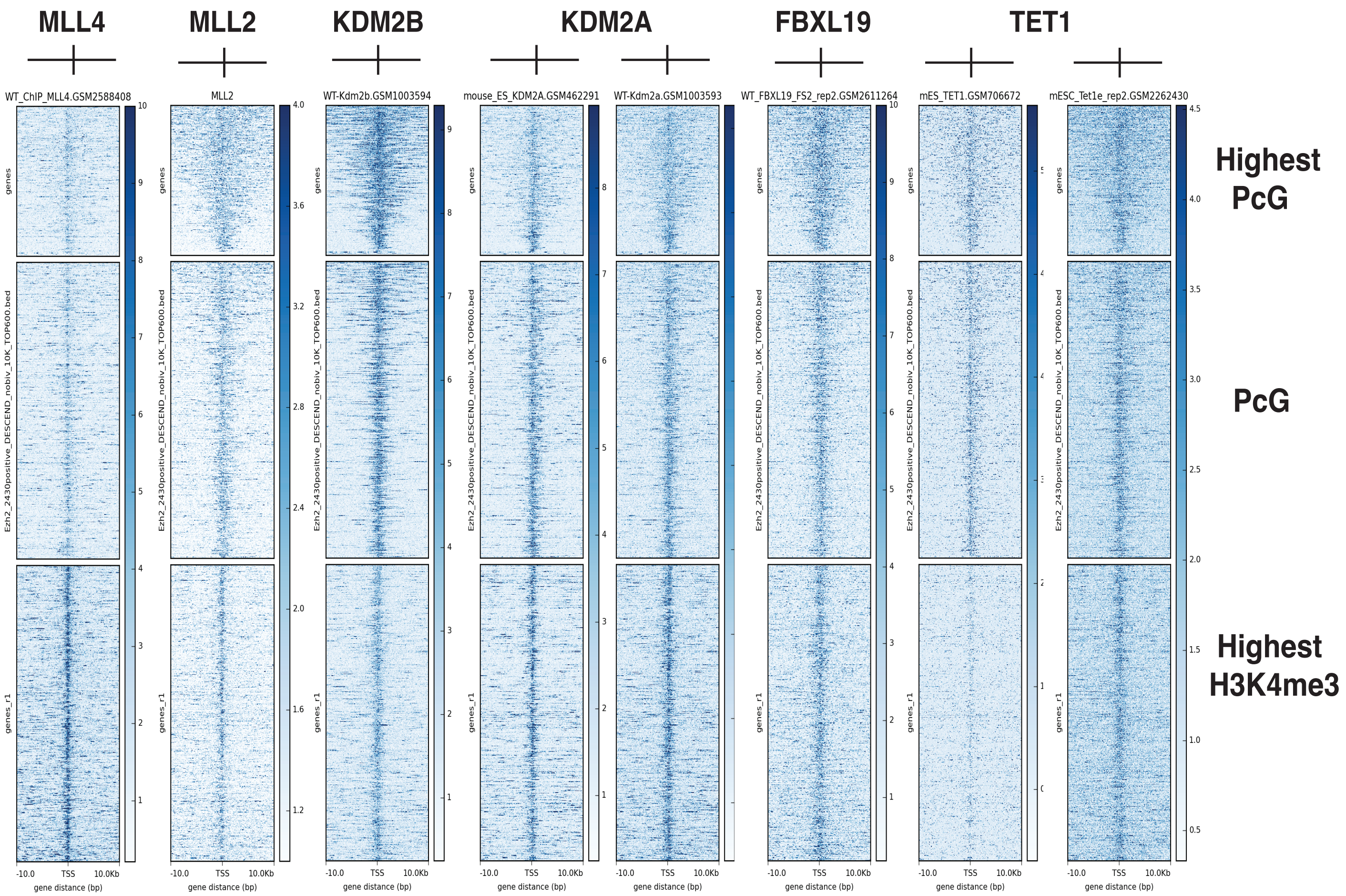
MLL4
MLL2
KDM2B
KDM2A
FBXL19
TET1
WT_ChIP_MLL4.GSM2588408
MLL2
WT-Kdm2b.GSM1003594
mouse_ES_KDM2A.GSM462291
WT-Kdm2a.GSM1003593
WT_FBXL19_FS2_rep2.GSM2611264
mES_TET1.GSM706672
mESC_Tet1e_rep2.GSM2262430
genes
Ezh2_2430positive_DESCEND_nobiv_10K_TOP600.bed
genes_r1
-10.0
TSS
10.0Kb
gene distance (bp)
Highest
PcG
PcG
Highest
H3K4me3

### Ji_FigS6

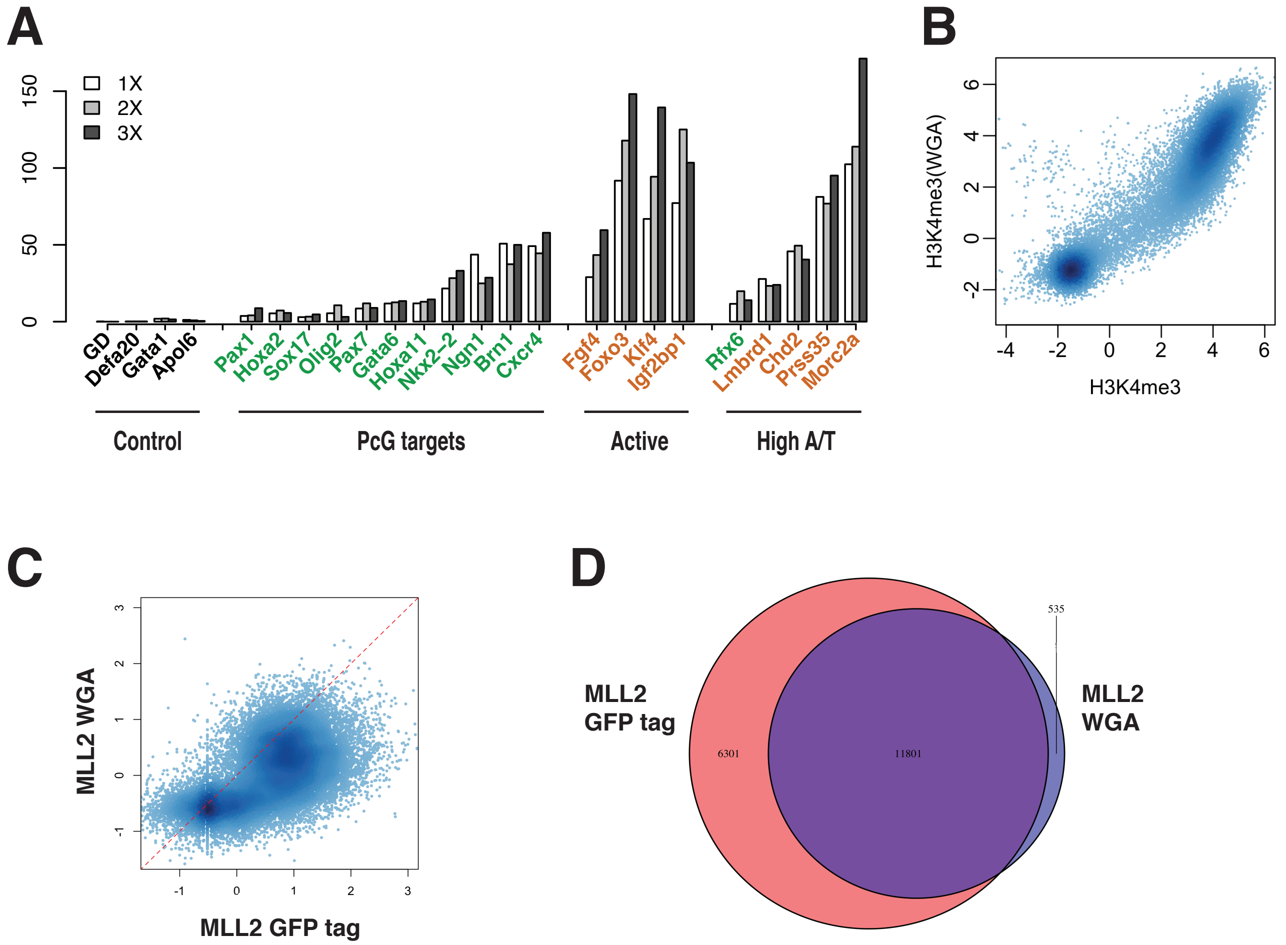
A
1X
2X
3X
0
50
100
150
GD
Defa20
Gata1
Apol6
Pax1
Hoxa2
Sox17
Olig2
Pax7
Gata6
Hoxa11
Nkx2-2
Ngn1
Brn1
Cxcr4
Fgf4
Foxo3
Klf4
Igf2bp1
Rfx6
Lmbrd1
Chd2
Prss35
Morc2a
Control
PcG targets
Active
High A/T
B
H3K4me3(WGA)
H3K4me3
-4
-2
0
2
4
6
C
MLL2 WGA
MLL2 GFP tag
-1
0
1
2
3
D
MLL2
GFP tag
MLL2
WGA
6301
11801
535

Ji_FigS7

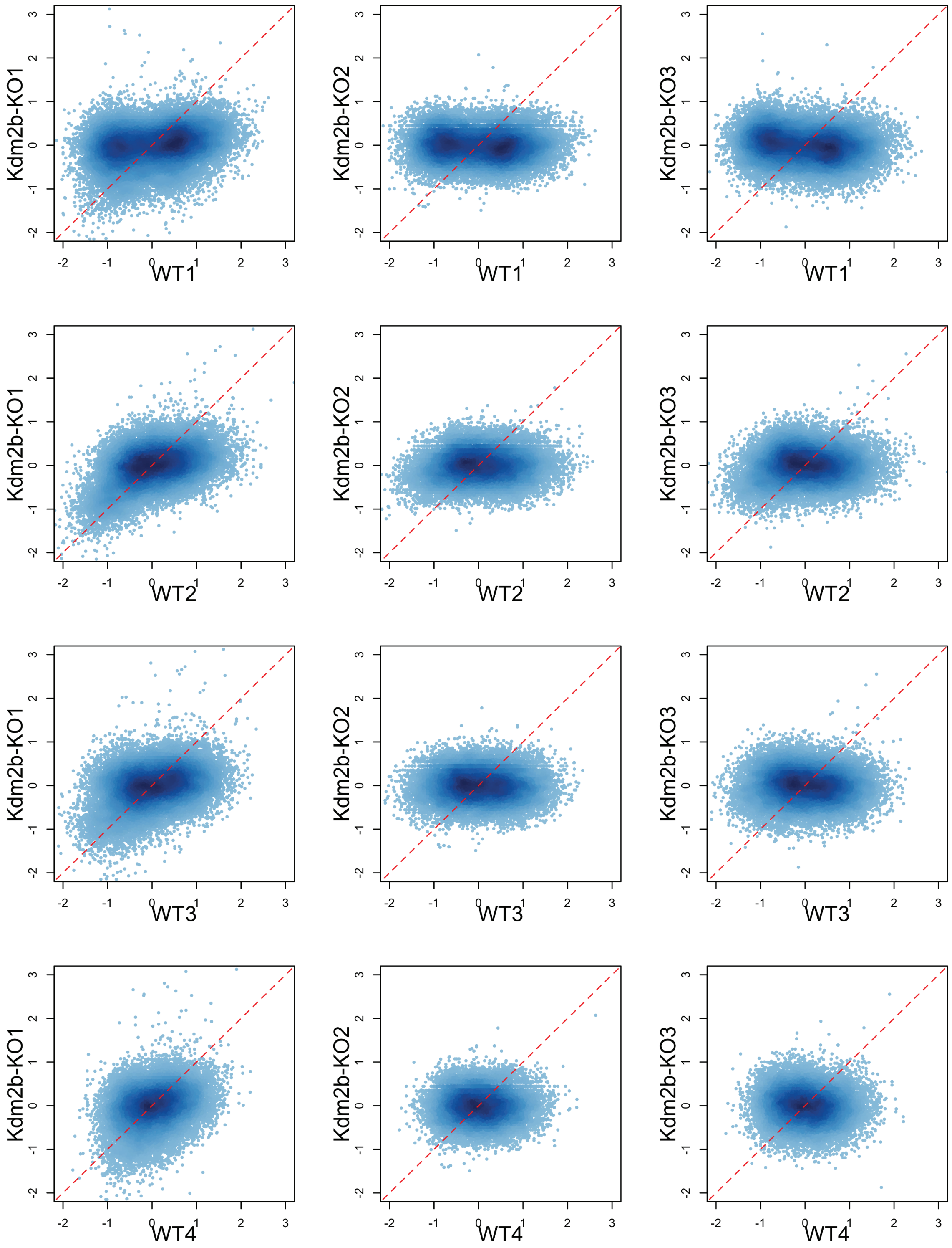

### Ji_FigS8

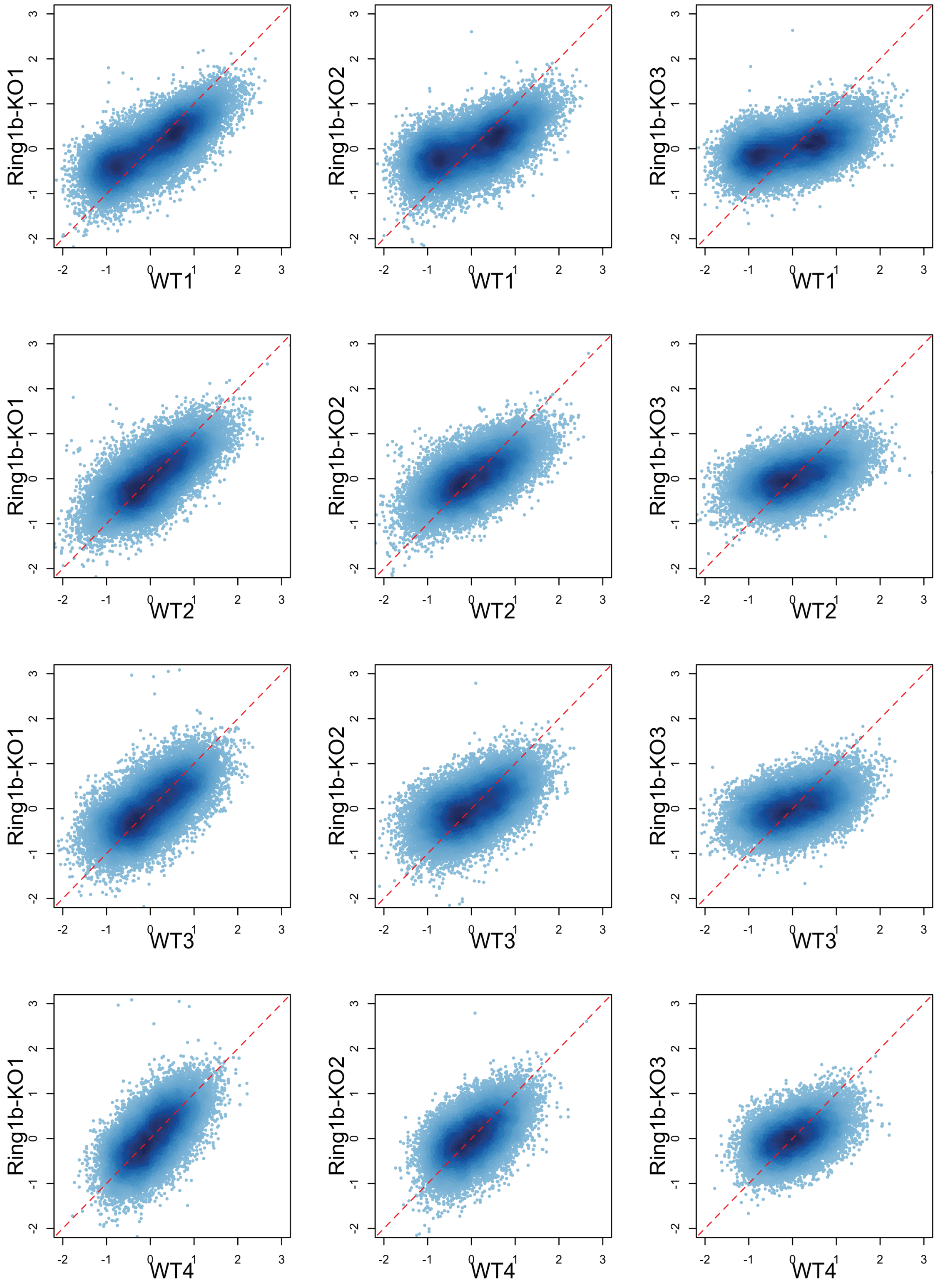

### Ji_FigS9

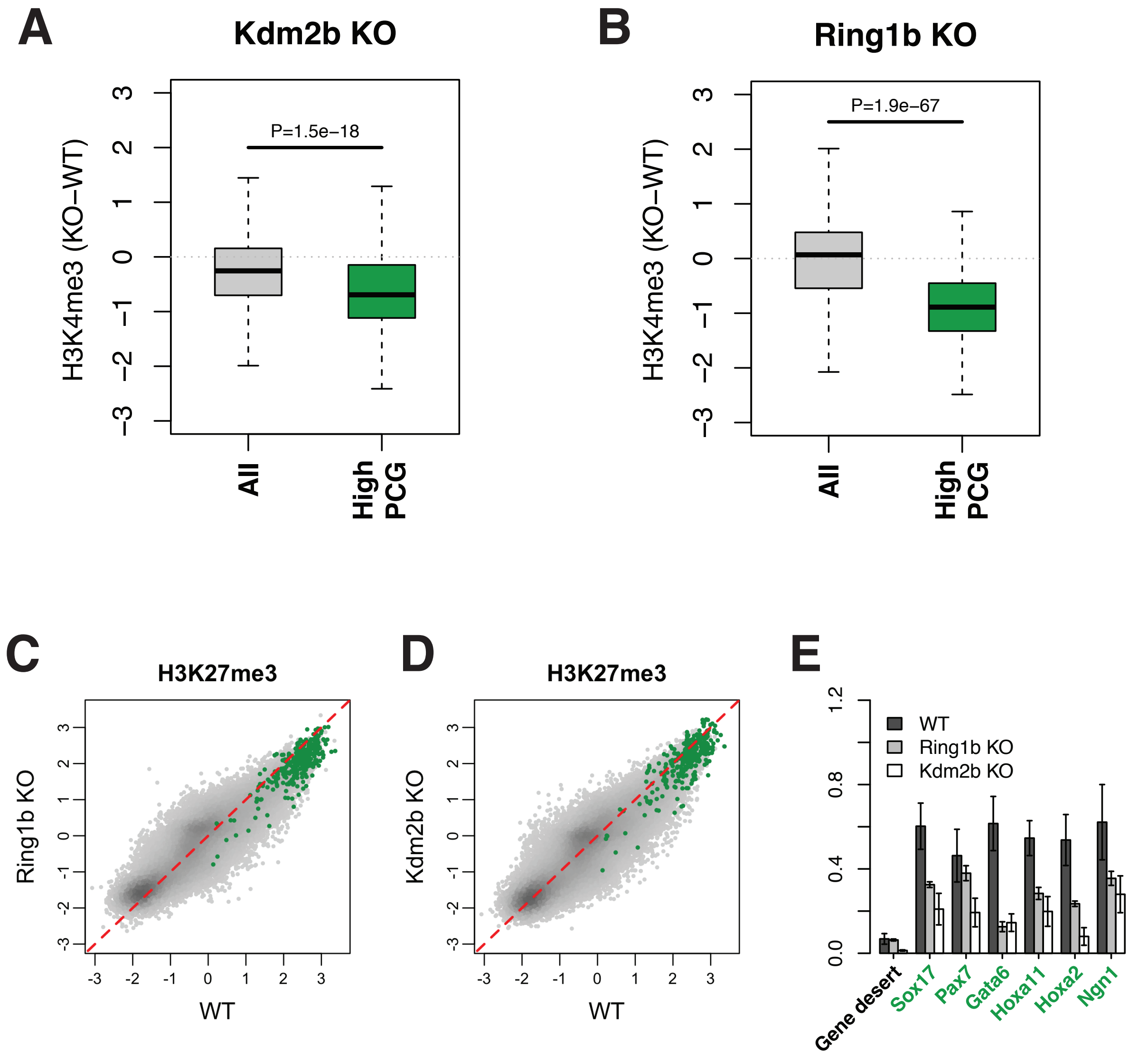

A
Kdm2b KO
H3K4me3 (KO−WT)
P=1.5e−18
All
High PCG
B
Ring1b KO
H3K4me3 (KO−WT)
P=1.9e−67
All
High PCG
C
H3K27me3
Ring1b KO
WT
D
H3K27me3
Kdm2b KO
WT
E
WT
Ring1b KO
Kdm2b KO
Gene desert
Sox17
Pax7
Gata6
Hoxa11
Hoxa2
Ngn1

### Ji_FigS10

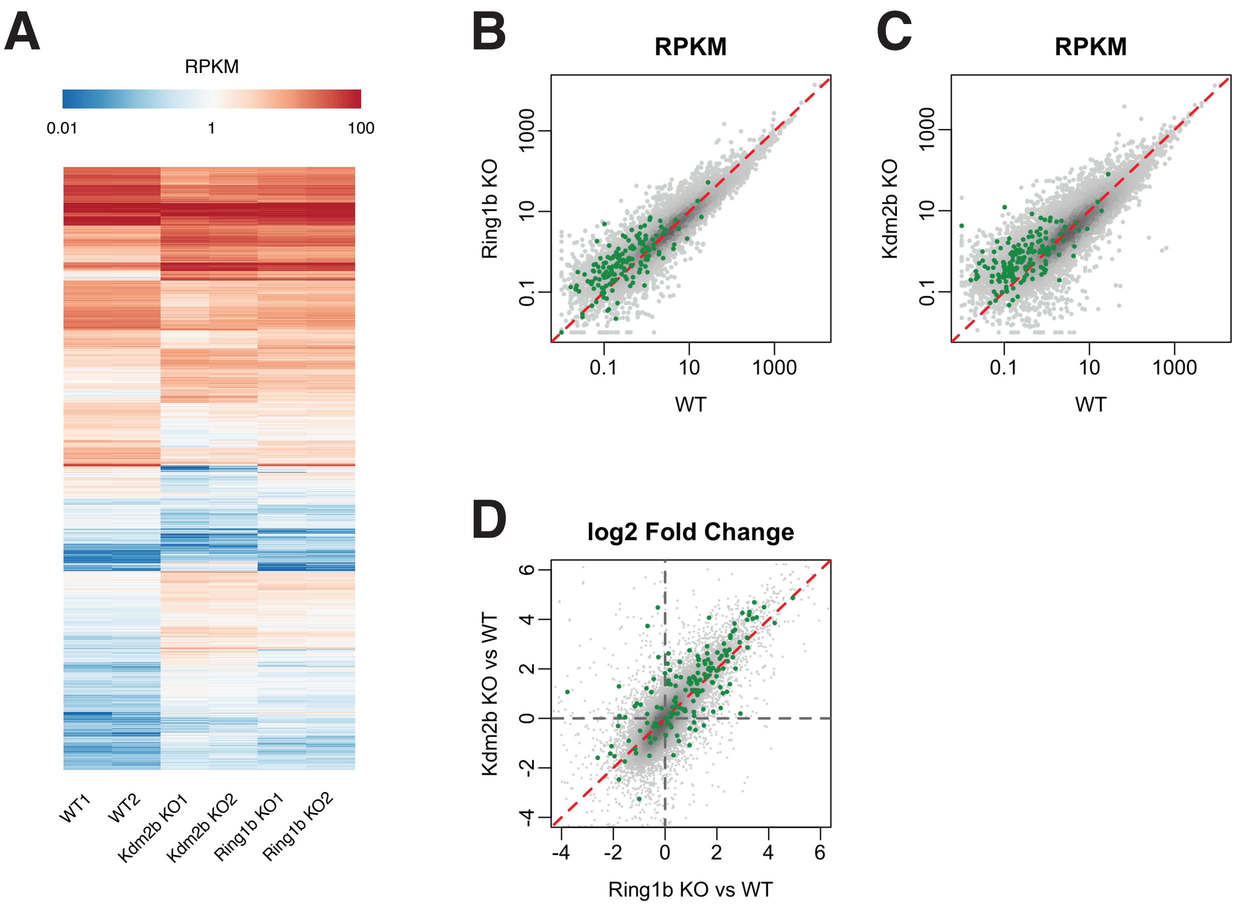
A
RPKM
0.01
1
100
WT1
WT2
Kdm2b KO1
Kdm2b KO2
Ring1b KO1
Ring1b KO2
B
RPKM
Ring1b KO
0.1
10
1000
WT
C
RPKM
Kdm2b KO
D
log2 Fold Change
Kdm2b KO vs WT
Ring1b KO vs WT
-4
-2
0
2
4
6
